# Pipeline for transferring annotations between proteins beyond globular domains

**DOI:** 10.1101/2022.11.08.515674

**Authors:** Elizabeth Martínez-Pérez, Mátyás Pajkos, Silvio C.E. Tosatto, Toby J. Gibson, Zsuzsanna Dosztanyi, Cristina Marino-Buslje

## Abstract

**Background:** DisProt is the primary repository of Intrinsically Disordered Proteins. This database is manually curated and the annotations there have strong experimental support. Currently DisProt contains a relatively small number of proteins highlighting the importance of transferring verified disorder and other annotations, in such a way as to increase the number of proteins that could benefit from this valuable information. While the principles and practicalities of homology transfer are well-established for globular proteins, these are largely lacking for disordered proteins.

**Methods:** We used DisProt to evaluate the transferability of the annotation terms to orthologous proteins. For each protein, we looked for their orthologs, with the assumption that they will have a similar function. Then, for each protein and their orthologs we made multiple sequence alignments (MSAs). Global and regional quality of the MSAs was evaluated with the NorMD score.

**Results:** We have designed a pipeline to obtain good quality MSAs and to transfer annotations from any protein to their orthologs. Applying the pipeline to DisProt proteins, from the 1931 entries with 5,623 annotations we can reach 97,555 orthologs and transfer a total of 301,190 terms by homology. We also provide a web server for consulting the results of DisProt proteins and execute the pipeline for any other protein. The server Homology Transfer IDP (HoTIDP) is accessible at http://hotidp.leloir.org.ar.

## INTRODUCTION

The structure of around 30% of the eukaryotic protein residues have never been determined by experimental techniques^1^, are inaccessible to template-based modelling and residues cannot be assigned to a Pfam family, a database of families of protein domains grouped by sequence similarity^2^. Most of these unmapped proteins or regions are predicted as disordered or compositionally biased according to several methods and databases such as IUPred3^3^, MobiDB^4^, AlphaFold^5^, and others.

The primary repository of disorder-related data of Intrinsically Disordered Proteins (IDPs) is DisProt^6^, a manually curated database, that ensures that each annotation has an experimental support. DisProt is the result of the effort of more than 60 experts. Disordered status is defined at a region level and the annotations are enriched with functional ontology terms. The Intrinsically Disordered Proteins Ontology (IDPO) collects structural and functional terms specific to IDPs and has been refactored and systematically cross-referenced with Gene Ontology (GO)^7^. Despite the great advantage of being manually curated, DisProt currently contains a relatively small number of proteins. This is because curating annotations in general, and for disordered proteins in particular, is a labour-intensive and time-consuming process, and that direct experimental evidence is also available for a limited number of proteins.

This highlights the importance of transferring annotations regarding verified disorder state and corresponding functions to homologous proteins, adding highly valuable information to better understand their biological roles. We focus on orthologous proteins as they are likely to have similar functions while paralogous proteins may or may not have similar functions. Homology transfer is well-established for globular proteins and is usually based on protein domain family annotations, such as PFAM^2^. However, the principles for homology transfer for Intrinsically Disordered Regions (IDRs), which often show larger evolutionary variation, is much less established.

A major problem to be faced when mapping features between homologous protein sequences is the variable quality of the sequences^8^. Only a small number of proteins have themselves been directly sequenced. The vast majority of sequenced DNA has been obtained through large scale genome sequencing initiatives of variable quality^9^. Any group of homologous sequences extracted from the protein sequence databases is likely to contain a mix of high and low quality entries.

Most multiple sequence alignment algorithms carry the assumption that the sequences to be aligned are collinear. When this condition is not met, alignment quality may be impacted^10^. If that happens, then misalignment may lead to errors in feature mapping. Therefore, in a sequence comparison pipeline, it makes sense to remove the most obviously problematic sequences early in the procedure. We expect the proteins in DisProt to be high quality since they are actively researched. Therefore they can act as references against which the problem proteins can be removed from the homology set. Generating high quality alignments will then allow annotation transfer with high confidence.

This work has two main goals, on the one hand, to provide to the community a protocol to safely transfer annotations from any protein to their orthologs. On the other hand, to transfer IDPO and GO annotations from DisProt proteins to their orthologs. Finally we’ve made a web server to bring the use of this protocol to the general public.

## MATERIALS AND METHODS

### DisProt database

We downloaded version 9.1 (2022-03) of the DisProt^6^ database. It currently collects regions in 2,365 proteins with a total of 6,763 manually curated IDPO terms and 3,251 GO terms. **Supplementary Figure 1** has an example of regions with GO and IDPO terms of protein TP53 potentially suitable to be transferred to its orthologs.

We filtered the dataset considering proteins following the rules: i) UniProt canonical proteins, ii) full length proteins, iii) sequences having no undefined amino acids (“X”) and iv) the protein sequence in DisProt and UniProt are identical. We ended up with 2,294 proteins, 3,156 GO terms and 6,550 IDPO terms annotated.

### Protein dataset

To avoid sequence redundancy, we clustered DisProt proteins with CD-HIT^11,12^, taking the longest one as the reference.

We collected the orthologs for every reference sequence from OmaDB^13^ and OrthoInspector^14^. We then added one-to-one orthologous proteins (the relationship between the pair of orthologs) for each reference sequence. Choosing one-to-one orthologs decreases the possibility of adding paralogs to the alignments. The sets of proteins obtained from the two databases were merged. Sequences having less than 30% coverage or more than 30% of the length of the reference one, were discarded. Also, all the clusters with only one sequence were removed.

### Multiple Sequence Alignments

Each cluster with their orthologs was aligned with Clustal Omega^15^ and MAFFT^16^. We also tested two sequence alignment conditions: sequences with less than 60% and 80% identity to the reference one (respectively) were removed from the MSAs. **Figure 1** shows an example MSA of Calreticulin (CALR) and its orthologs where removing sequences with less than 60% identity to the reference, changes a bad (**Figure 1A**) into a good MSA (**Figure 1B**) as measured with NorMD.

**Figure 1.**
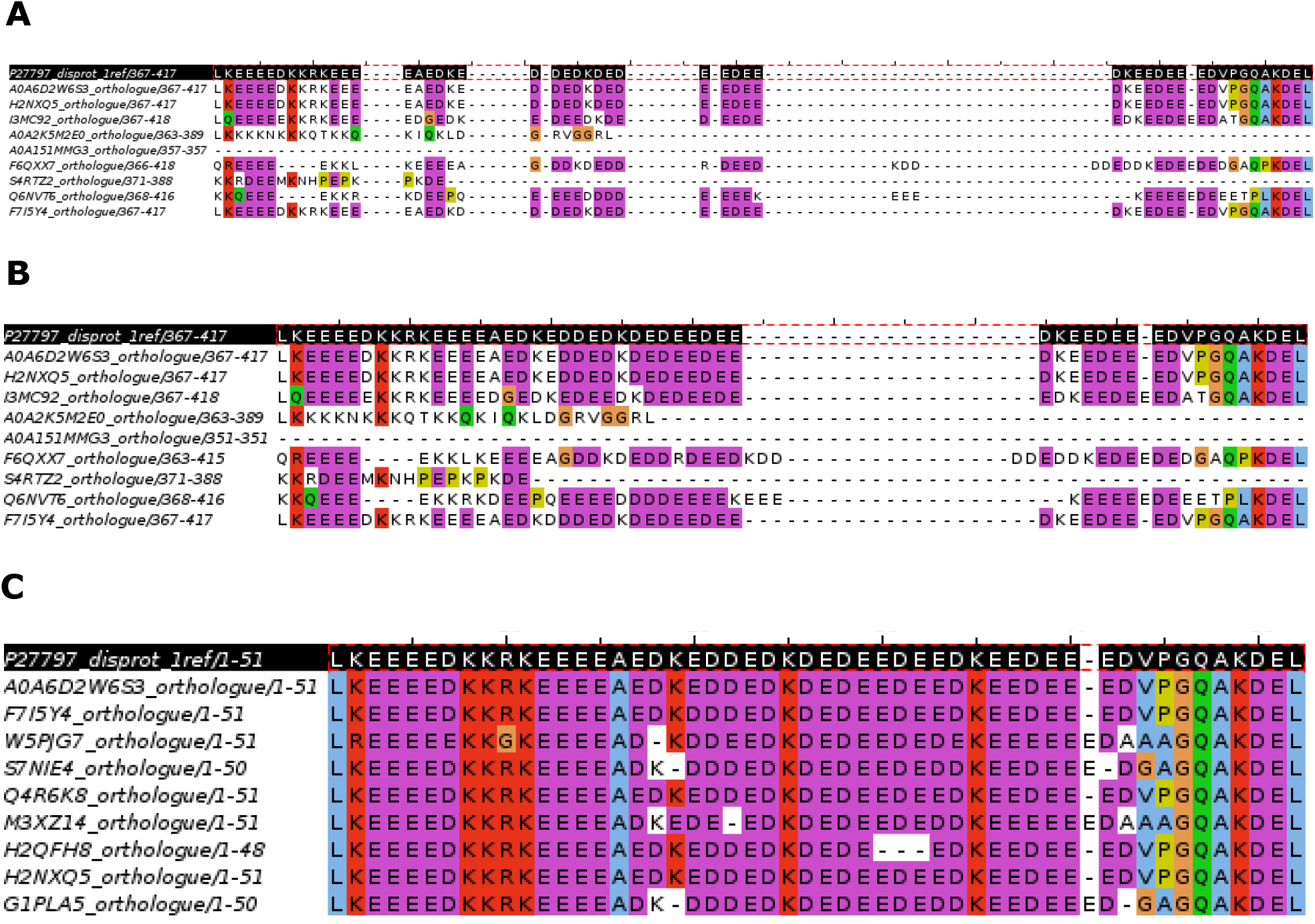
Example of the MSA of CALR, UniProt accession P27797, and DisProt ID DP00333. The region containing residues 367-417 is annotated in DisProt with the term “disorder” (IDPO:00076) (black row at top of the MSA). Only 10 sequences of the full alignment are shown so gap-only columns are present in this figure **A:** global MSA with 254 sequences has a bad NorMD score (0.496). **B:** global MSA after eliminating sequences less than 60% identity to the reference one (170 sequences), has a good global NorMD score (0.848) but this particular region still has a bad NorMD score (0.251). **C:** region (DisProt ID DP00333r007) MSA after eliminating sequences less than 60% identity in this region (50 sequences) now has a good NorMD score (1.000). MSA images were processed with Jalview^19^.

Then we chopped the global alignment and aligned each region containing a term separately. Here also, we tested removing sequences with less than 60% and 80% identity to the reference one in the aligned region. **Figure 1C** shows the annotated region (IDPO:00076) aligned after removing sequences with less than 60% identity in the region.

We evaluated the quality of each MSA with the NorMD^17^ score. The advantage of the NorMD score is the combination of the column-scoring and the residue similarity scores. Additionally, NorMD includes *ab initio* sequence information, like the amount, length and similarity of the sequences to be aligned. So, the NorMD score gives information about the general quality of the alignment. A NorMD score >0.6 is considered to indicate a reliable MSA^18^.

Note that proteins in DisProt can have overlapping annotations which might differ partially because region boundaries sometimes are defined by the construct used in the corresponding experiment. Therefore, the alignment corresponding to each annotation is evaluated separately. The exact boundary of the regions can influence which sequences are removed from the alignment of a particular region and influence which terms are transferred to other sequences.

### Testing homology transfer within DisProt

We took advantage of the fact that some alignments have more than one DisProt protein (the reference and non-reference ones) with annotated GO and IDPO region terms. Using these other members, we tested whether the annotation that we transferred to the non-reference sequence overlaps with the actual one annotated in the non-reference proteins. The overlap of a given region was calculated as the percentage of aligned amino acids annotated with a GO or IDPO term between the reference and the non-reference protein region. In the cases where the aligned region of a given non-reference protein is not annotated in DisProt, the overlap is zero, even if the identity between the regions is very high (these cases fall into the 0-10 bin). The same applies at different % overlap for regions that are not annotated in exactly the same positions between the 2 proteins (**Supplementary Figure 2** represents the different situations).

### WorkFlow chart

We developed a workflow to test the quality of alignments and consequently, the reliability of annotation transfer (**Supplementary Figure 3)** shows the general WorkFlow Chart used in this work. Briefly, we took the DisProt entries and clustered them. In parallel, we looked for their orthologous proteins. We then aligned each cluster including their orthologs with Clustal Omega and filtered the sequences based on the sequence identity to the reference one, to avoid redundancy. We considered two quality scores: one corresponding to the global alignment (full protein length) and the other to the annotated regions with GO or IDPO terms such as: disorder to order transition, protein binding, etc (an example of terms are in **Supplementary Figure 1**).

### Data and codes accessibility

All the data and codes are available at the IDPfun GitLab project https://gitlab.com/idpfun/homology_transfer_disprot. Web Server: Homology Transfer (HoTIDP) is accessible at http://hotidp.leloir.org.ar.

## RESULTS

DisProt^6^ is the reference database of Intrinsically Disordered Proteins. In the database, disordered regions are enriched with structural and/or functional annotations. In order to test the usability of its information through homology transfer to other proteins, we collected orthologous proteins from two databases, OrthoInspector^11^ and OmaDB^12^. From the 2,294 proteins in the DisProt database only 1,931 have available orthologs (**Supplementary Table 1**). A total of 579,647 one-to-one orthologous proteins were retrieved from the two databases (**Figure 2A**). As the ortholog databases are different, reference proteins might have orthologs in one database and not in the other. Also the number of orthologs for a protein in each database can vary. This highlights the importance of using more than one orthology database. Merging them allows us to increase the number of orthologs available to potentially extend the annotations via homology transfer.

**Figure 2.**
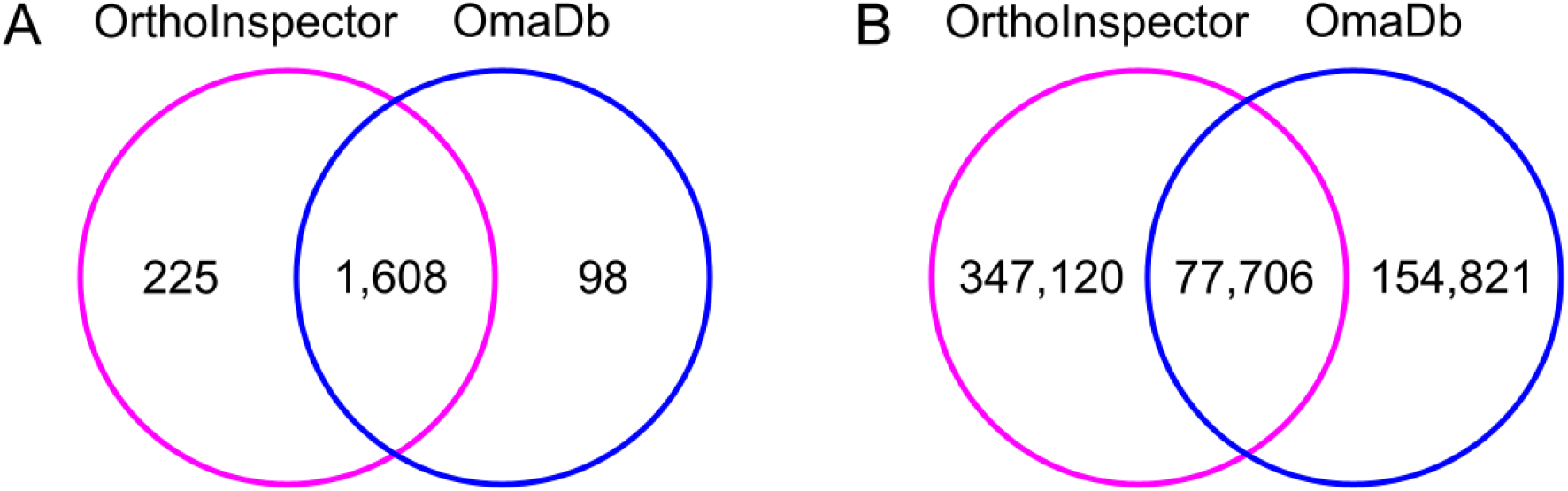
Datasets of orthologous proteins. **A:** Venn diagram of DisProt proteins having one-to-one orthologs in each database. **B:** Venn diagram of one-to-one orthologs retrieved from DisProt proteins.

However, a key requirement for a reliable homology transfer is a good quality MSA. As disorder and corresponding functional annotations are in general assigned to particular segments of the protein, it is also important to examine alignment quality not just at the global level, but also at the level of regions.

### Comparison between different parameters

We analysed different parameters for assigning the suitability of an annotation term to be transferred. The MSAs with each set of parameters and the number of proteins, are included in **Table 1**. The NorMD score was computed for each MSA, a distribution is shown in the **Supplementary Figure 4**.

**Table 1.** Numbers for the 60% and 80% datasets.

| Initial DisProt proteins %identity Clustering | Alignment method | %identity Global MSA | Global MSAs | Number of proteins | %identity Region MSA | Region MSAs | Number of proteins |
| --- | --- | --- | --- | --- | --- | --- | --- |
| 60% | Clustal Omega | >= 60% | 1,728 | 128,664 | >= 60% | 3,295 | 96,571 |
|  |  |  |  |  | >= 80% | 3,128 | 70,799 |
|  |  | >= 80% | 1,621 | 61,460 | >= 60% | 3,157 | 57,865 |
|  |  |  |  |  | >= 80% | 3,064 | 52,077 |
|  | Mafft | >= 60% | 1,730 | 134,111 | >= 60% | 3,304 | 100,753 |
|  |  |  |  |  | >= 80% | 3,130 | 72,879 |
|  |  | >= 80% | 1,628 | 63,684 | >= 60% | 3,166 | 60,089 |
|  |  |  |  |  | >= 80% | 3,070 | 53,923 |
| 80% | Clustal Omega | >= 60% | 1,776 | 131,033 | >= 60% | 3,342 | 99,286 |
|  |  |  |  |  | >= 80% | 3,184 | 72,311 |
|  |  | >= 80% | 1,674 | 63,476 | >= 60% | 3,231 | 59,578 |
|  |  |  |  |  | >= 80% | 3,135 | 53,604 |
|  | Mafft | >= 60% | 1,778 | 137,084 | >= 60% | 3,352 | 104,250 |
|  |  |  |  |  | >= 80% | 3,188 | 74,504 |
|  |  | >= 80% | 1,682 | 65,740 | >= 60% | 3,241 | 61,833 |
|  |  |  |  |  | >= 80% | 3,143 | 55,484 |
| Total |  |  | 13,617 | 137,528 |  | 51,086 | 104,909 |

Highlighted in bold is the line of work explained in the manuscript for clarity, although all these possibilities were tested and results are exposed in the supplementary material.

Statistical differences were found between the NorMD score of the MSAs generated by different sets of parameters (Kruskal-Wallis test, p-value 1.537349e-222 < 0.05). To analyse the differences, we made pairwise comparisons between the set of parameters highlighted in **Table 1** with the other sets using Dunn’s test corrected by False Discovery Rate (see **Supplementary Table 2)**. As expected, alignments having more than 80% pairwise identity, are more accurate based on the NorMD score than alignments at 60% pairwise identity (p-values <0.05) and there is no dependence neither with the initial DisProt percent identity clustering nor with the alignment method (p-values >0.05).

It is worth noting that all the global and region alignments with sequences >60% pairwise identity have a good NorMD score (>0.6).

### Suggested annotation transferring for DisProt proteins

If the global alignment has a good score while having a bad score in the annotated region and vice versa, these terms are not suitable to be transferred.

Our initial dataset contained 2,294 proteins with 3,156 GO terms and 6,550 IDPO terms annotated. These proteins were clustered at 80% identity with CD-HIT^11,12^, ending up with 2,151 clusters. Out of these, a total of 1,931 sequences had a total of 579,647 one-to-one orthologs. After basic quality checks, we ended up with 1,849 clusters with more than one entry. For each cluster, we generated multiple sequence alignment with Clustal Omega.

After filtering, a total of 97,555 proteins could be assigned with 301,190 homology transferred terms (84,380 are GO, and 220,886 are IDPO terms).

### Testing homology transfer within DisProt

In total, 222 alignments had more than one protein from DisProt. In 156 cases (70%), the percentage of overlap between annotations was at least 1%. For 61% of the cases (136 cases) the overlap was more than 50%, and 54% of the cases (119 cases) the overlap was better than 80%.

However 66 cases (30%) have no overlap between the annotations. The overlap between regions has a double-peaked distribution (shown in **Figure 3**). The absence of overlap might be because many regions are not annotated for all the DisProt entries, albeit having high identity with the reference protein.

**Figure 3.**
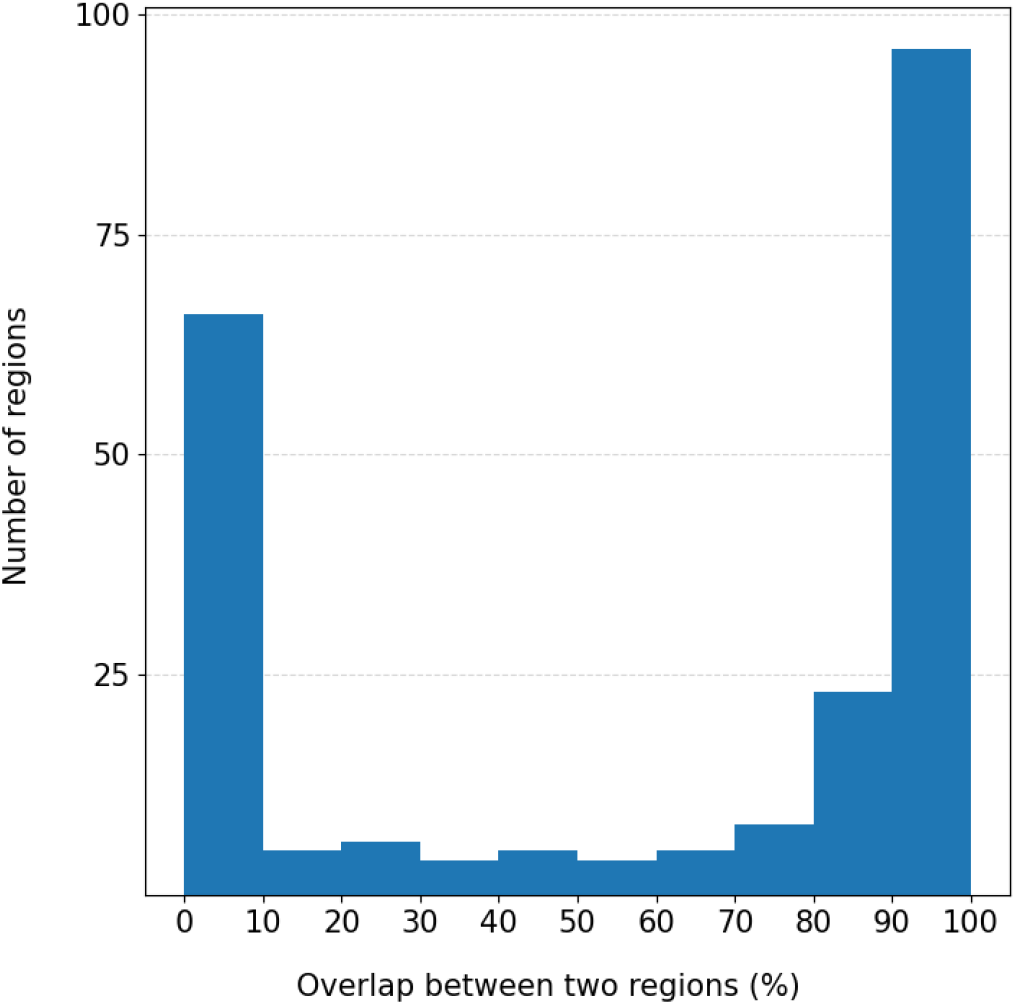
Distribution of overlapped regions between reference and non-reference proteins.

### Pipeline to transfer terms by homology

The pipeline for generating high quality MSAs, and transferring the region annotations is available at https://gitlab.com/idpfun/homology_transfer_disprot. The software requirements are programming languages (Python, R and Perl), workflow control (snakemake), alignment methods (Clustal Omega and MAFFT), and measuring the MSA quality (norMD score). The pipeline can be downloaded and executed on a personal computer. The default parameters are described in this manuscript, and can also be modified in the snake configuration file.

### Server HoTIDP Homology Transfer Database

We developed a web server HoTIDP that provides an interface and programmatic access to obtain the transferable terms if a protein has orthologs in DisProt.

The server can be queried with a protein UniProt accession and, if it has an ortholog in DisProt, the annotated regions and transferable terms will be provided. Also it shows the protein family alignment as well as a pairwise alignment with the reference protein. A table with all the comparison scores (alignment quality of the different regions, and identity % with the reference protein) is also provided.

The default pipeline uses 80% DisProt clustering, Clustal Omega and global and local alignment at 60% of identity to the reference protein. However, the user can select any of the tested parameters (initial clustering, aligning methods and the global and local % identity)

It can also suggest transferring a term to any protein if a reference protein and region is provided by the user. The procedure is making an MSA with good quality global and local alignment scores with the orthologous proteins. The server is hosted in http://hotidp.leloir.org.ar.

## DISCUSSION

In this work we delineated guidelines to transfer annotations to orthologous proteins. In such a way to enrich their functional and structural information, in particular for intrinsically disordered proteins. We collected 579,647 one-to-one orthologs from 1,931 DisProt proteins. We aligned each family of proteins with Clustal Omega and stored the MSAs that surpass a quality threshold. We compared MSAs at different conditions: clustering DisProt proteins by 60% and 80% identity, keeping orthologous sequences 60% and 80% identity to the reference protein in the global and region MSAs, and two alignment methods. Our results show that the quality of the alignment does not depend on the alignment method for this similarity cutoffs. As expected, the quality of MSAs constructed gathering sequences more than 80% identical to the reference one, are significantly different (and better) to MSAs made with sequences with more than 60% of identity. However, all MSAs with sequences more than 60% identity also have a good NorMD score.

The MSAs having more than one DisProt protein allowed us to test the method on 222 regions, 136 of which have an overlap of at least 50% of their residues with the reference protein (**Figure 3**). The overlap between reference and non-reference protein regions may be underestimated due to differences in the annotation of the two DisProt entries. For instance, the human, mouse and rat Epsin orthologs are annotated in DisProt, however the term at human residues 1 to 18 is not annotated in the closely related orthologs, although these proteins are nearly 100% identical, probably because there is no experimental evidence to support it. This annotation gives important insights into the protein functions, such as that it is disordered, has a transition to an ordered state, and also is a small molecule sequestering region. However, the current available data limits us to do an exhaustive validation.

Sequences having good NorMD scores in the global alignment (even with high sequence identity) can still have regions with low quality scores. If we omit checking the quality of the alignments in the regions, one third of the annotations (414,025 terms to 129,257 proteins), would be transferred incorrectly. After filtering sequences in each annotated region of the MSAs we safely transferred 301,190 terms to 97,555 proteins. This highlights the importance of having this protocol instead of transferring, like other methods do, just considering the identity of the global alignment^4^.

This protocol increases by ∼40 times the amount of disordered proteins with functional annotations, and ∼25 times the annotated regions with GO terms and ∼32 times with IDPO terms. As an example, **Table 2** shows the regions that GO and IDPO terms could be transferred from the reference FUS protein to the orthologous proteins.

**TABLE 2.** Example of annotated regions that could be transferred from human FUS protein to the orthologs.

| Start | End | MSA proteins | Term ID | Term name |
| --- | --- | --- | --- | --- |
| 1 | 163 | 49 | GO:0140693 | molecular condensate scaffold activity |
|  |  |  | GO:0043232 | intracellular non-membrane-bounded organelle |
|  |  |  | IDPO:00076 | disorder |
| 1 | 214 | 45 | GO:0043232 | intracellular non-membrane-bounded organelle |
|  |  |  | GO:1990000 | amyloid fibril formation |
| 1 | 422 | 50 | GO:0043232 | intracellular non-membrane-bounded organelle |
| 1 | 507 | 52 | IDPO:00076 | disorder |
| 1 | 526 | 54 | GO:0140693 | molecular condensate scaffold activity |
|  |  |  | GO:0043232 | intracellular non-membrane-bounded organelle |
| 2 | 36 | 49 | IDPO:00076 | disorder |
| 37 | 41 | 53 | GO:0005515 | protein binding |
| 39 | 95 | 49 | GO:1990000 | amyloid fibril formation |
| 98 | 214 | 43 | IDPO:00076 | disorder |
| 149 | 154 | 48 | GO:0005515 | protein binding |
| 285 | 370 | 53 | GO:0003723 | RNA binding |
|  |  |  | IDPO:00050 | disorder to order |
| 421 | 455 | 53 | IDPO:00050 | disorder to order |
| 508 | 526 | 45 | GO:0005515 | protein binding |

FUS is taken as the canonical sequence (UniProt accession P35637)

In summary, we developed guidelines to transfer terms that might give relevant information to homolog proteins.

We also developed a web server HoTIDP that can be used to transfer any term to any protein if the UniProt accession and the regions with the terms are provided.

When we apply the pipeline to the DisProt proteins the GO and IDPO terms annotated can be reliably transferred to more than 97,000 proteins increasing their information.

## ACKNOWLEDGEMENTS

We thank Dr. Julie D. Thompson who kindly provided us with the NorMD software and helped with the setting up.

## FUNDING

- This work is part of a project that has received funding from the European Union’s Horizon 2020 research and innovation programme under the Marie Skłodowska-Curie grant agreement No. 778247” OR “Supported by the H2020-MSCA-RISE project IDPfun-GA No. 778247.”
- EMP is a Postdoctoral fellow and CMB is a researcher of the Argentine National Research Council (CONICET).

## Supplementaries

**Supplementary Material: Supplementary Figures 1, 2, 3 and 4 and Supplementary Tables 1 and 2**

https://docs.google.com/document/d/1V3wLUNo-Oi3Igj9tqL0A9_XrW9bHOWjLUMlIvBOAld0/edit?usp=sharing

## Notes

http://hotidp.leloir.org.ar

### Competing Interest Statement

The authors have declared no competing interest.

