## Supplementary Material for "Pipeline for transferring annotations between proteins beyond globular domains"

#### Supplementary Figure 1.

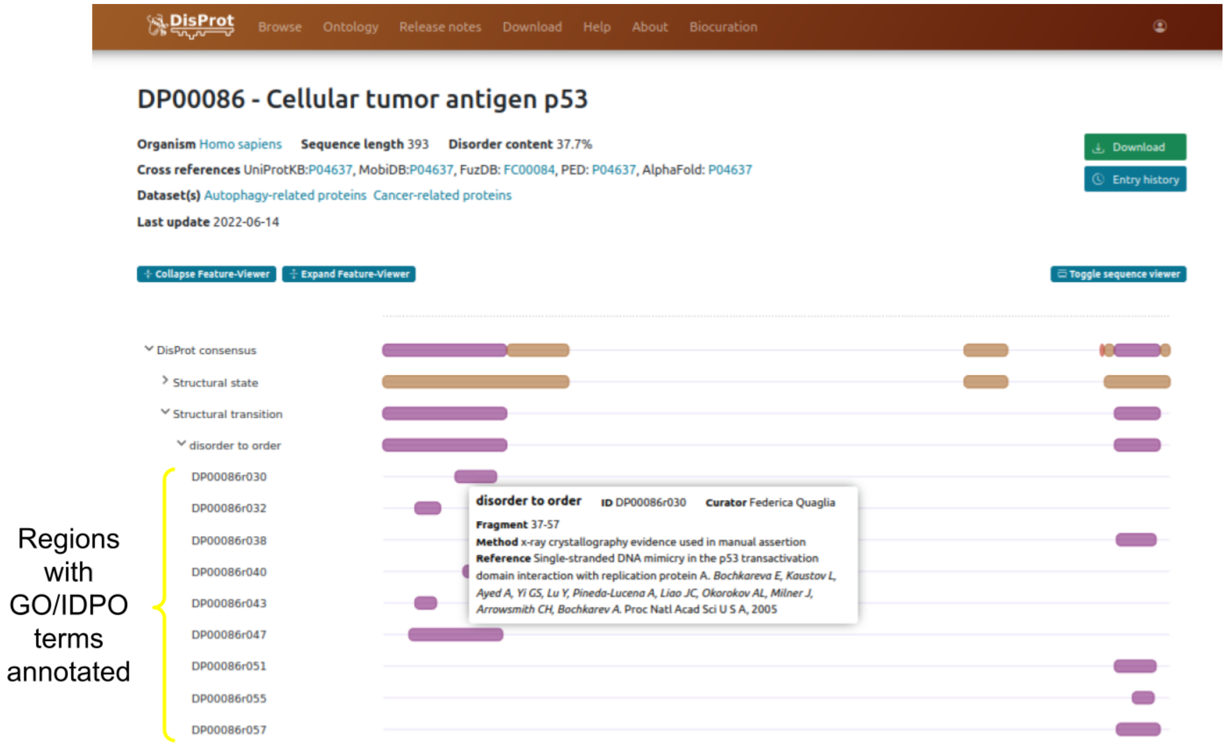


**Supplementary Figure 1. Example of regions with GO/IDPO terms for P53.** Screenshot showing detail for DisProt accession DP00086. At the top level of the ontology are the Structural state, Structural transition and Disordered function. The other terms are in the second level of the ontology, and the notations like DP00086r030 are regions with the specific IDPO terms.

### Supplementary Figure 2.

```
       Reference: MASNDYTQQATQSYGAYPTQPGQGYSQQSSQPYGQQSYSGYSQSTDTSGYGQSSYSSYGQ0
A 100% overlap: MASNDYTQQATQSYGAYPTQPGQGYSQQSSQPYGQQSYSGYSQSTDTSGYGQSSYSSYGQ0
B 100% overlap: MASNDYTQQATQSYGAYPTQPGQGYSQQSSQPYGQQSYSGYSQSTDTSGYGQSSYSSYGQ0
C  50% overlap: MASNDYTQQATQSYGAYPTQPGQGYSQQSSQPYGQQSYSGYSQSTDTSGYGQSSYSSYGQ0
D  50% overlap: M-----------------TQPGQGYSQQSSQPYGQQSYSGYSQSTDTSGYGQSSYSSYGQ0
E   0% overlap: MASNDYTQQATQSYGAYPTQPGQGYSQQSSQPYGQQSYSGYSQSTDTSGYGQSSYSSYGQ0
F   0% overlap: M-----------------------------------SYSGYSQSTDTSGYGQSSYSSYGQ0
```

**Supplementary Figure 2. Examples of aligned IDR sequence regions.** The Reference sequence is FUS (UniProt Accession P35637) amino acids 1-60. The orange box shows the range of the annotated "disorder" (term IDPO:00076) for amino acids 2 to 36. The sequences A-F and their annotations are artificial examples. 0% overlap means no annotation of this region in the other DisProt protein (sequence E) or a region not present in the orthologue (Sequence F).

**Supplementary Figure 3.**

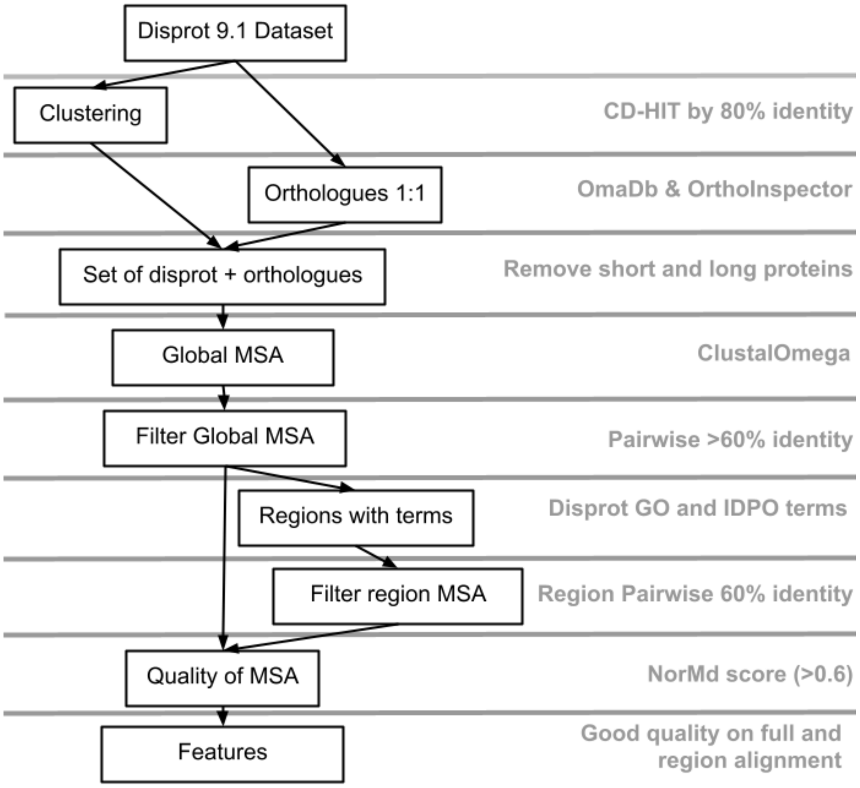


**Figure 3.** Outline of the workflow. Reference DisProt sequences are used to collect orthologs and produce MSAs. After quality evaluation, annotated features are transferred.

### Supplementary Figure 4.

A

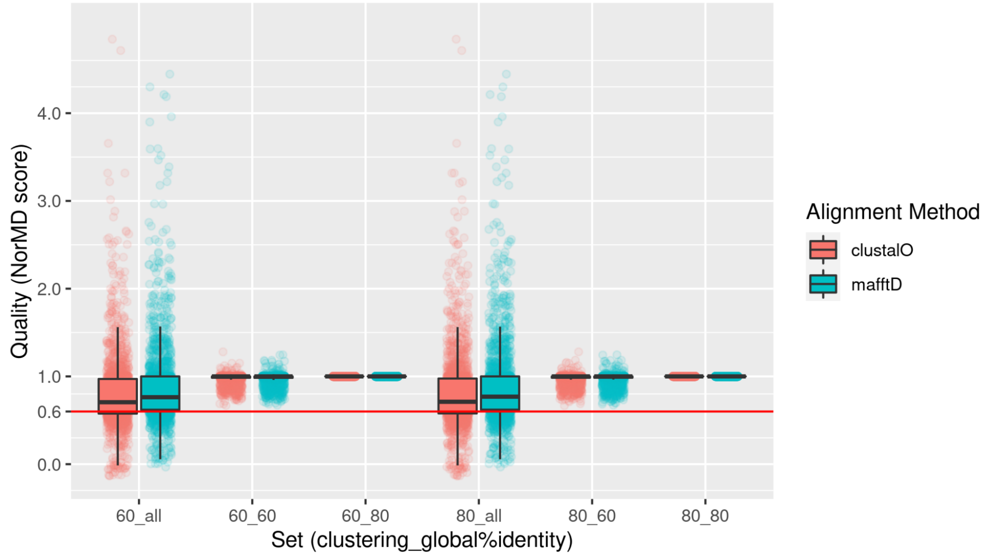


B

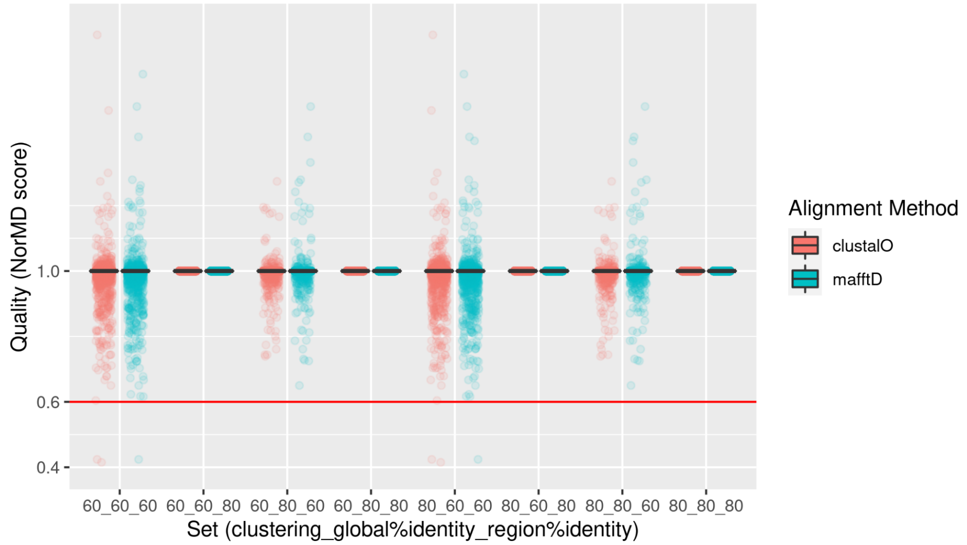


**Supplementary Figure 4. NorMD scores for MSAs. A: NorMD score in global MSAs. B: NorMD score in region MSAs.** The "y" axis is the NorMD score. The "x" axis are the sets to compare using the nomenclature AA_BB or AA_BB_CC where AA is clustering type, BB the % on the global MSAs and CC is the %identity on the region MSAs. All the %identity mentioned are between each sequence and the reference in the MSA. The BB as "all" are global MSA without filter sequence by %identity.

**Supplementary Table 1. DisProt proteins and their orthologs found in the databases OmaDB and OrthoInspector.**

| | **OmaDB** | **OrthoInspector** | **Total** |
|---|---|---|---|
| DisProt proteins not found | 509 | 429 | |
| DisProt proteins found with valid one-to-one orthologs | 1,697 | 1,827 | 1,924 |
| Valid orthologs found | 230,997 | 424,030 | 577,653 |

We consider valid orthologues when the protein is: not obsolete in UniProt, has a Complete sequence (no fragments), and has no "X" in the primary sequence.

**Supplementary Table 2.** Comparison the NorMD scores between sets of alignments, using the Dunn test and the p-value are corrected with FDR.

| Disprot clustering | Alignment method | Global %identity+ | Region %identity+ | P-value comparing with Default parameters* |
|---|---|---|---|---|
| 60 | Clustal Omega | 60 | 60 | **8.39E-01** |
| 60 | Clustal Omega | 60 | 80 | 3.41E-43 |
| 80 | Clustal Omega | 60 | 80 | 3.41E-43 |
| 60 | Clustal Omega | 80 | 60 | 1.91E-15 |
| 60 | Clustal Omega | 80 | 80 | 7.16E-43 |
| 80 | Clustal Omega | 80 | 60 | 1.89E-15 |
| 80 | Clustal Omega | 80 | 80 | 3.41E-43 |
| 60 | MAFFT | 60 | 60 | **4.50E-01** |
| 60 | MAFFT | 60 | 80 | 3.41E-43 |
| 80 | MAFFT | 60 | 60 | **4.50E-01** |
| 80 | MAFFT | 60 | 80 | 3.41E-43 |
| 60 | MAFFT | 80 | 60 | 2.28E-18 |
| 60 | MAFFT | 80 | 80 | 7.16E-43 |
| 80 | MAFFT | 80 | 60 | 7.85E-19 |
| 80 | MAFFT | 80 | 80 | 3.41E-43 |

**+ %identity** is the pairwise value between each sequence and the reference sequence

*** Default parameter** is DisProt clustering 80% identity, Clustal Omega as alignment method, 60% identity in global and region Pairwise alignment with the reference protein.
